# Learning the histone codes of gene regulation with large genomic windows and three-dimensional chromatin interactions using transformer

**DOI:** 10.1101/2021.12.30.472333

**Authors:** Dohoon Lee, Jeewon Yang, Sun Kim

## Abstract

The quantitative characterization of the transcriptional control by histone modifications (HMs) has been challenged by many computational studies, but still most of them exploit only partial aspects of intricate mechanisms involved in gene regulation, leaving a room for improvement. We present Chromoformer, a new transformer-based deep learning architecture that achieves the state-of-the-art performance in the quantitative deciphering of the histone codes of gene regulation. The core essence of Chromoformer architecture lies in the three variants of attention operation, each specialized to model individual hierarchy of three-dimensional (3D) transcriptional regulation including (1) histone codes at core promoters, (2) pairwise interaction between a core promoter and a distal *cis*-regulatory element mediated by 3D chromatin interactions, and (3) the collective effect of the pairwise *cis*-regulations. In-depth interpretation of the trained model behavior based on attention scores suggests that Chromoformer adaptively exploits the distant dependencies between HMs associated with transcription initiation and elongation. We also demonstrate that the quantitative kinetics of transcription factories and polycomb group bodies, in which the coordinated gene regulation occurs through spatial sequestration of genes with regulatory elements, can be captured by Chromoformer. Together, our study shows the great power of attention-based deep learning as a versatile modeling approach for the complex epigenetic landscape of gene regulation and highlights its potential as an effective toolkit that facilitates scientific discoveries in computational epigenetics.

## Introduction

The control of gene expression is carried out by diverse groups of regulators including transcription factors, coactivators, corepressors along with the genomic sequence elements. However, the basic premise behind the interplay among those factors is the appropriate configuration of the covalent modifications of histone tails, or histone modifications (HMs), at the relevant genomic regions since they play a pivotal role in the regulation of the chromatin accessibility. Thus, it can be conceived that the amount of HMs and their combinations encode the regulatory potential of the nearby genomic regions. This notion is referred to as the ‘histone code hypothesis’^1^. There have been a number of computational and quantitative approaches to crack the regulatory code of gene expression encoded by HMs. Most of them are predictive models that utilize the levels of HMs at promoters surrounding transcription start sites (TSSs) to predict the expression level of the corresponding gene. Notably, recent studies have shown the superior performance of deep learning models compared to the conventional machine learning models in this task^2, 3^.

To date, deep learning has been making remarkable breakthroughs in diverse fields of computational biology, ranging from the characterization of binding specificity of DNA- and RNA-binding proteins^4^ to the longstanding problem of the protein structure prediction based on its amino acid sequence^5^. These successes of deep learning in biology could not be achieved without the invention of novel model architectures and also their clever applications for complex biological problems. In that sense, the high complexity of histone code indeed made it a great target for deep learning as shown in the existing approaches, but they still pose two major limitations that motivate the development of a new approach.

One is that they could only use narrow genomic windows around TSSs. This is because the deep learning architectures that those models were based on, such as convolutional neural networks (CNNs) and recurrent neural networks (RNNs), were not effective in modeling the dependencies within long sequences. CNNs are highly specialized for learning local patterns of data, but it is challenging for them to learn the distant dependencies between the patterns. Although developed to model sequential data, RNN architectures also have difficulties in capturing the long-range dependencies clearly since the information embedded in a single position becomes gradually diluted and gets contaminated while the model computation travels along the positions between the two distant positions. Although advanced forms of RNN cells such as Gated Recurrent Units^6^ or Long Short-Term Memory (LSTM)^7^ partially ameliorates this problem, the intrinsic inefficiency in modeling of long sequences due to the recurrence still remains.

Next, a majority of the deep learning models do not account for the distal *cis*-regulation mediated by three-dimensional (3D) chromatin folding, even though it has been widely known that the physical interactions between core promoters and distal *cis*-regulatory elements critically modulates the gene expression^8, 9^. In other words, the regulatory information conveyed by the histone code is allowed to not only propagate locally, but also jump between distant genomic loci through 3D chromatin interactions^10^. Fortunately, the recent advancement of high-throughput measurement technologies such as Hi-C^11^ succeeded in providing a fine-resolution view of 3D chromatin interactions at kilobase-scale and offered us with unprecedented opportunities for exploiting such valuable information to model the comprehensive view of gene regulation. There are few emerging studies that explicitly take the 3D chromatin interactions into consideration to predict the gene expression. One such example is GC-MERGE^12^, a graph neural network (GNN) to propagate information between interacting genomic regions to predict the expression levels of genes. Although it is a proof-of-concept model that cannot be applied to genes without any chromatin interactions and only performs 10kbp genomic bin-level predictions but not at gene-level, it still underscores the promise of modeling epigenomic contexts of distal genomic regions along with that of promoters.

In an effort to overcome these limitations, we present a novel transformer-based deep learning architecture named Chromoformer to predict the gene expression levels based on the HMs at the wide neighborhood of TSSs as well as the HMs placed at the distal regulatory elements. The transformer architecture was originally developed for natural language processing^13^, but recently it has also been exhibiting great potential for understanding the latent grammar of DNA sequences^14^, amino acid sequences^15^ and even their alignments^16^. In particular, in this study we noticed that the two main functionalities of the transformer architecture are highly suitable to tackle the two aforementioned challenges. First, transformers can precisely model the long-range dependencies in sequential data. This is elegantly done by the addition of positional encodings to input sequences. These input features harboring positional information are treated independently and fed into a subsequent self-attention module which calculates the all-pairwise dependencies between the input features. Therefore, long-range dependencies can be captured without the interference of features located between the pair. Secondly, the transformer architecture can also be applied to model unordered sets of entities along with the interactions among them. Of note, this is not straightforward for most of the deep learning architectures since the operations comprising them depends on input positions. On the other hand, the operations comprising the transformer are basically permutation-invariant. The interactions between input features are only considered in self-attention operations and all the other operations are done in position-wise manner, so it can be applied to model unordered set of features.

Together, these two strengths of the transformer architecture make it a promising choice for the quantitative modeling of histone codes by allowing us to utilize wider genomic windows near TSSs and histone codes at multiple distal regulatory regions simultaneously. By exploiting these advantages of transformers, we designed a novel deep learning model architecture consisting of three variants of self-attention operations to reflect the hierarchy of 3D gene regulation. As a result, Chromoformer model achieved far better predictive power compared to the other deep learning models for gene expression prediction. Moreover, through the comprehensive investigation on how the use of transformer architecture contributed to the superior performance of the model, we could demonstrate that the long-range modeling of epigenetic context near TSS and simultaneous integrative modeling of distal regulatory regions actually worked to improve performances. Finally, we showed that we could draw artificial intelligence-driven hypotheses for the quantitative effect of *cis*-regulation by the two subdomains within nuclei, transcription factories and silencing hubs, through the interpretation of the dynamics of latent embeddings of the regulatory states learned by Chromoformer.

## Results

### Chromoformer adopts three-level transformer architecture that reflects the hierarchy of 3D gene regulation

The core design principle of Chromoformer is twofold. One is to extract as much proximal regulatory information as possible from the HMs at the core promoters, and the other is to incorporate the distant histone codes whose information is transmitted to the core promoter through 3D chromatin interactions. To fully utilize the transformer architecture to model the complex dynamics of *cis*-regulations involving multiple layers, we conceptually decomposed the gene regulation into a three-layered hierarchy: (1) *cis*-regulation by core promoters, (2) 3D pairwise interaction between a core promoter and a putative *cis*-regulatory regions (pCREs) and (3) a collective regulatory effect imposed by the set of 3D pairwise interactions. To computationally emulate this hierarchy, we introduced three transformer-based submodules called Embedding, Pairwise Interaction and Regulation transformers that are specialized to learn the respective grammar of gene expression regulation in the order of increasing complexity.

Before illustrating the model architecture, we briefly describe the input features used throughout this study. Chromoformer was trained using read depth values from histone ChIP-seq experiments for seven major HMs (H3K4me1, H3K4me3, H3K9me3, H3K27me3, H3K36me3, H3K27ac and H3K9ac). Read depths were averaged and log2-transformed for fixed-sized bins across 40kbp regions flanking TSSs (Figure 1a). To account for the distal *cis*-regulation, we additionally utilized the HM signals at pCREs that are known to interact with the core promoter in the corresponding cell type. For that, an experimentally validated set of pCREs for each core promoter was obtained using a publicly available collection of promoter-capture Hi-C (pcHi-C) data^17^. The 3D chromatin interactions were characterized at the resolution of HindIII restriction fragments. The interactions were characterized at adequately high-resolution, as the median and average length of those fragments were 4,797bp and 5,640bp, respectively and about 95% of them were less than 10kbp (Supplementary Figure 1).

**Figure 1.**
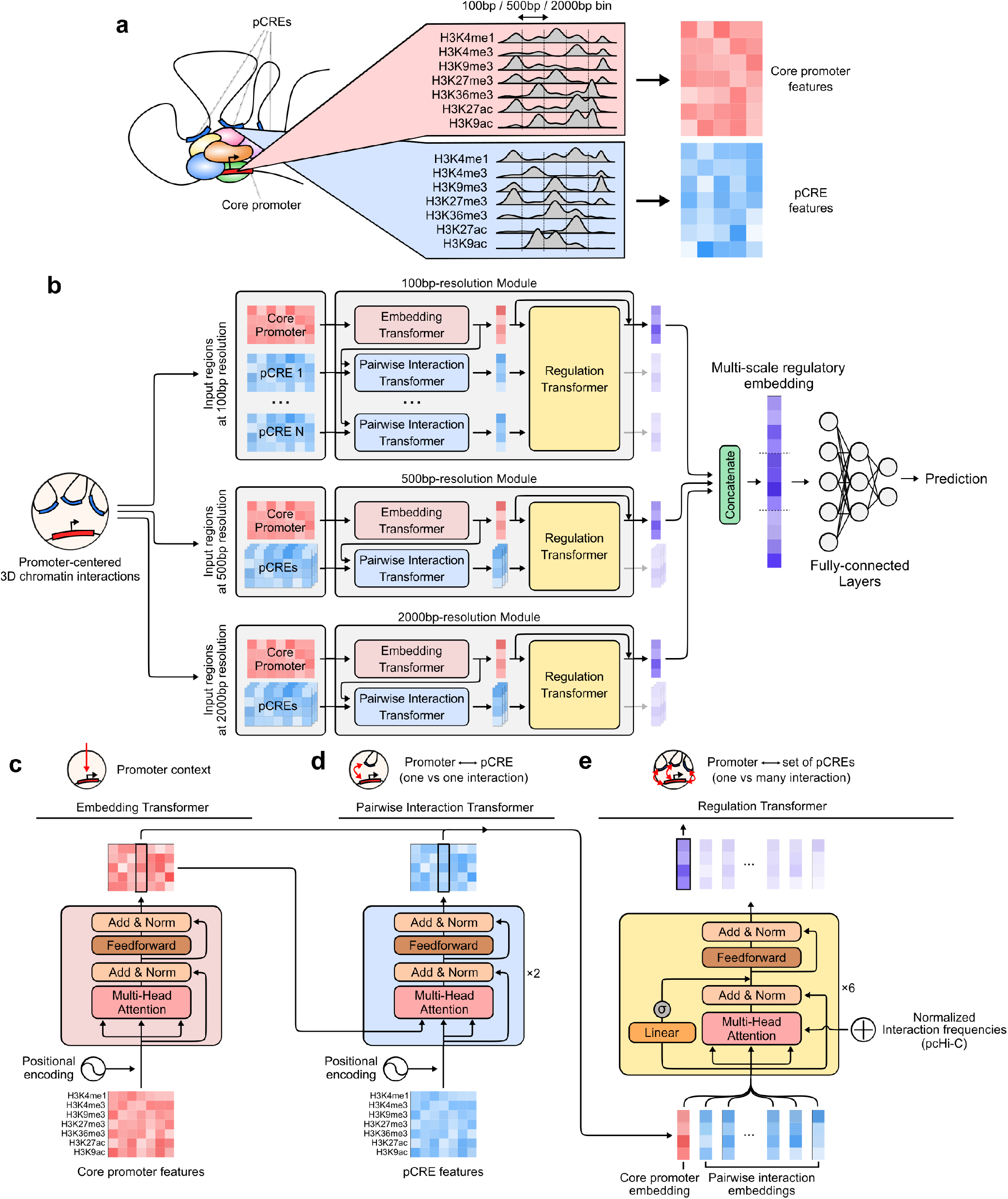
Chromoformer model architecture. (a) Input features. To predict the expression of a gene using levels of histone modifications (HMs), we extracted binned average signals of HMs from both the core promoter and putative *cis*-regulatory regions (pCREs). (b) Chromoformer architecture. Three independent modules were used to produce multi-scale representation of gene expression regulation. Each of the modules is fed with input HM features at different resolutions to produce an embedding vector reflecting the regulatory state of the core promoter. (c) Embedding transformer architecture. Position-encoded HM signals of core promoter features are transformed into core promoter embeddings through self-attention. (d) Pairwise Interaction transformer architecture. Position-encoded HM signals of pCREs are used to transform the core promoter embeddings into Pairwise Interaction embeddings through encoder-decoder attention. (e) Regulation transformer architecture. Using the whole set of the core promoter and Pairwise Interaction embeddings and gated self-attention, Regulation transformer learns how the pCREs collectively regulate the core promoter. To guide the model to put greater attention to frequently occurring three-dimensional (3D) interactions, the normalized interaction frequency vector is added to self-attention affinity matrices.

The full model architecture used in this study is illustrated in Figure 1b. At the highest level it consists of three independent modules each of which accepts input features at different resolutions and in turn produces an embedding vector of the regulatory state at the core promoter. The resulting three regulatory embeddings are concatenated to form a multi-scale regulatory embedding which is subsequently fed into fully-connected layers to predict the expression level of the gene. The Embedding transformer (Figure 1c) learns the histone codes acting at the direct vicinity of a TSS and produces a fixed-sized vector that summarizes the epigenetic state of the region. This submodule alone works very similar to the existing machine learning models for HM-based gene expression prediction, but we expected that the use of transformer architecture will allow the model to precisely identify relevant signals within a wide-range view (up to 40kbp) of core promoter HM contexts without any performance degradation. Next, the resulting core promoter embeddings are further updated by the Pairwise Interaction transformer (Figure 1d) in the context of pairwise *cis*-regulatory interactions between promoters and pCREs. Instead of using typical self-attention layers as in the Embedding transformer, this module is built with encoder-decoder attention layers. Since the activity of a promoter is modulated by the contact with pCREs, the encoder-decoder framework was chosen to reflect this by decoding the promoter embeddings given the context of the pCRE features. We call the resulting embedding vectors pairwise interaction embeddings as they carry the information of one-to-one relationship between promoters and pCREs. Finally, the Regulation transformer (Figure 1e) accepts a union set of the core promoter and pairwise interaction embeddings and finally produces a regulatory embedding by integrating them. This module models the whole landscape of *cis*-regulation using gated self-attention layers. The normalized interaction frequencies are injected to self-attention score matrices to guide the model with the priorities of interactions. Detailed explanation of the model is illustrated in Methods.

### Chromoformer outperforms existing deep models for gene expression prediction based on epigenetic features

We benchmarked the performance of Chromoformer with three baseline deep learning models using the optimal settings proposed in the respective studies. We first trained DeepChrome^2^, a convolutional neural network that learns the local combination of HMs through the weights of convolutional filters to predict gene expressions. We also trained the AttentiveChrome^3^ models that combines LSTM and global attention mechanism to enhance the interpretability of the model. Lastly, an HM-based hybrid CNN-RNN model proposed by Kang et al.^18^ (HM-CRNN) was also chosen for the comparison. It learns and captures the meaningful local combination of HMs through CNN and comprehends their sequential dependencies through RNN.

To measure the modeling performance, we let the models solve a task of gene expression prediction in a reduced form as a binary classification problem to predict whether an expression of a gene is above median or not. This problem formulation was first proposed by Singh et al.^2^, and it has been so far widely adopted for many studies including the three aforementioned baseline studies. Classification performances were evaluated for 11 cell types among the 127 cell types profiled by Roadmap Epigenomics^19^ and ENCODE project^20^. Those 11 cell types were chosen because all of the gene expression profiles, ChIP-seq data for seven major HMs and pcHi-C interaction profiles were publicly available for each of the cell types. A total of 18,955 genes were split into four sets for 4-fold cross-validation (CV), each consisting of 5,045, 4,751, 4,605 and 4,554 genes. For every CV fold, each set became a held-out validation set while the other three sets were used for model training. To avoid unwanted information leakage from training to validation set through 3D chromatin folding involving promoter-promoter interaction, we ensured that no two genes in different sets are located on the same chromosome.

As a result, our model based on multi-scale Chromoformer model showed significant performance improvement over existing baseline deep learning models in terms of area under receiver operating characteristic curve (ROC-AUC), suggesting that the transformer-based modeling of the regulatory hierarchy was successfully working (Figure 1a). Similarly, we observed consistent performance increase when average precision or accuracy were used as performance measures (Supplementary Figure 2a and b). These results were consistently reproduced for all 11 cell types examined. Besides, we found that the prediction probabilities produced by Chromoformer showed remarkable positive correlation with the actual expression levels (Supplementary Figure 2c). These well-calibrated prediction probabilities for the quantitative expression levels justifies the use of binary classification formulation for the quantitative modeling of HMs.

Furthermore, Chromoformer also far outperformed GC-MERGE, a GNN using three-dimensional chromatin interaction to predict gene expression (Figure 1b). Importantly, GC-MERGE can only predict for genes involved in at least one chromatin interaction. Also, GC-MERGE can only predict the gene expression in the unit of 10kbp genomic bins, therefore it cannot produce gene-wise prediction when two or more genes are present in the same bin. Therefore, Chromoformer was retrained from scratch for a subset of genes whose expression can be predicted by GC-MERGE for a fair comparison.

### Training with large window size and *cis*-regulatory interactions contributed to the performance improvement in Chromoformer

To dissect the performance of Chromoformer into the contributions of individual factors, we first inspected for the effect of modeling wide-range windows around the TSS up to 40kbp. By gradually increasing the window size around TSS from 2kbp to 40kbp, we observed a consistent performance increase for our model, while other deep learning models showed considerable performance degradation when the window size was larger than 10kbp (Figure 3a). Interestingly, widening the input window size seemed to be slightly more detrimental to the two attention-based baseline models, AttentiveChrome and HM-CRNN, than DeepChrome. We speculated that the global attention mechanism used in those models, which utilizes a few global context vectors to compute attention scores^21^, fails to put effective attention to the relevant signals when the window size is large. The large window size are more likely to include TSSs of other genes, which jeopardizes the model performance since the model will be more prone to the spurious attention towards those irrelevant TSSs. This is because the attention score for each genomic position is computed against global context vectors without accounting for the absolute distance to the target TSS. In the case of the transformer architecture, however, a scaled-dot product attention with positional encoding allows the computation of attention scores among all pairs of positions without introducing a global context vector, thereby allowing us to query on which position within the input window the specific TSS of our interest is attending on. Thus, these results emphasize the strength of the transformer architecture in pinpointing histone codes that are relevant to the regulation of gene expression within a wide-range window around the TSS.

**Figure 2.**
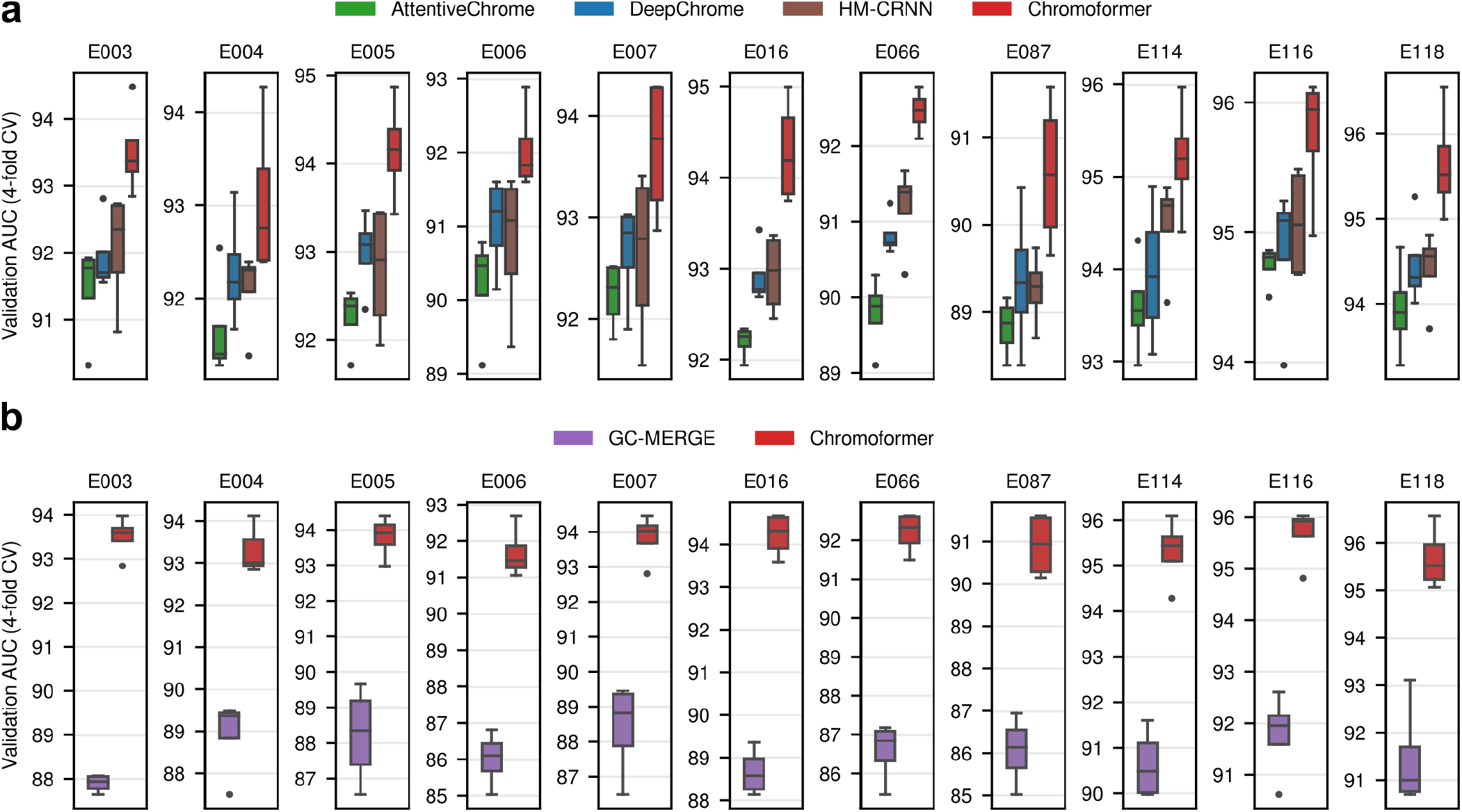
Chromoformer outperformed existing deep learning models in predicting gene expression states. (a) Performance comparison with the benchmark deep learning models that only utilize the core promoter features. (b) Comparison with GC-MERGE, a graph neural network model that utilizes 3D *cis*-regulatory interactions. For fair comparisons, Chromoformer models were retrained from scratch only using a subset of genes that GC-MERGE can predict. The boxes denote the interquartile range, and the whiskers denote the 1.5× interquartile range. AUC, Area under receiver operating characteristic curve.

**Figure 3.**
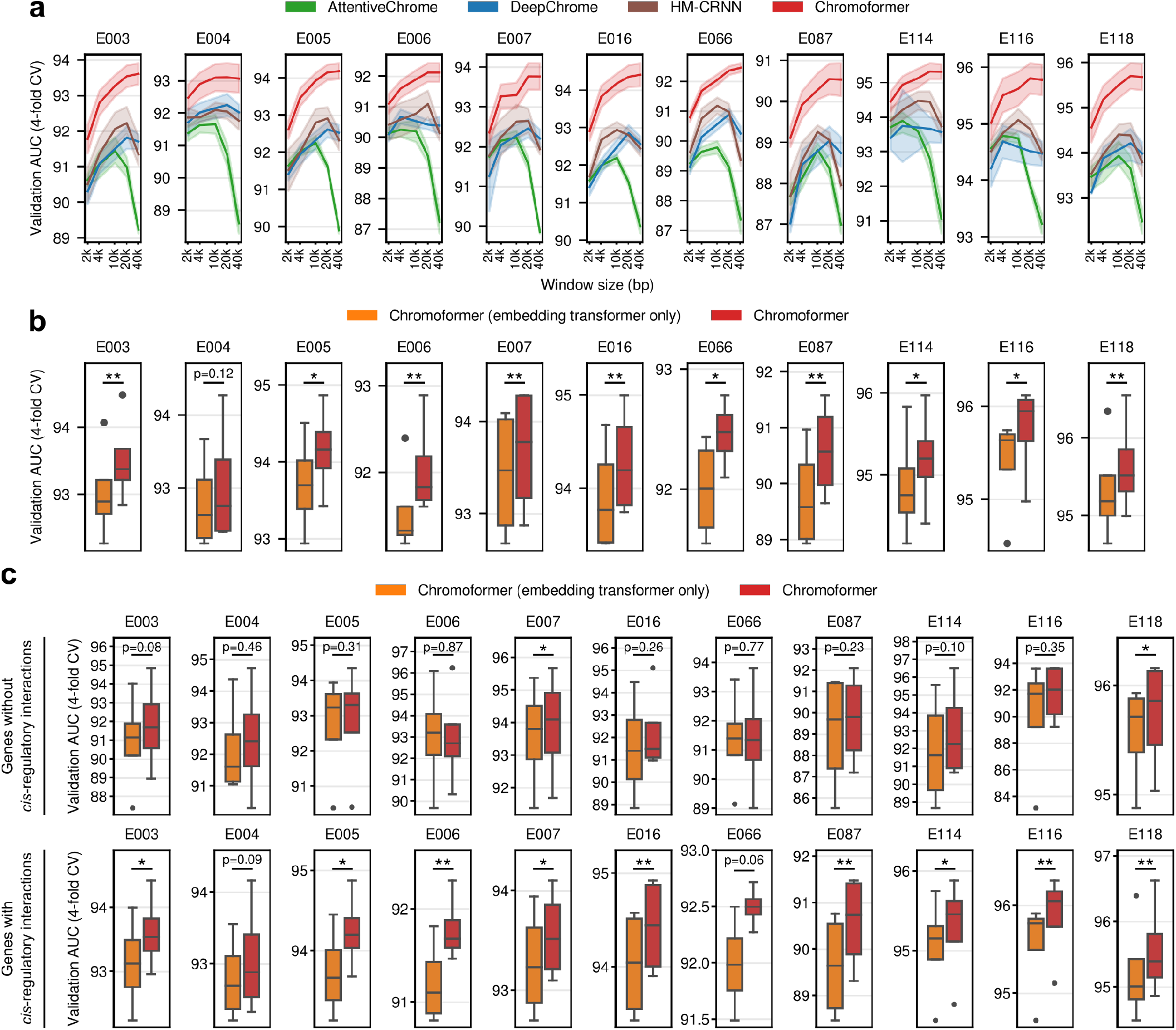
Contributing factors to the superior performance of Chromoformer. (a) Effect of input window size around TSS on the model performance. Chromoformer and the other benchmark models were trained for five different window sizes (2kbp, 4kbp, 10kbp, 20kbp and 40kbp), while all the other training procedures were kept the same as previously. The shades denote the standard error across 4-fold cross validation for each window size. (b) Effect of taking distal *cis*-regulations by pCREs into account. We trained ablated Chromoformer models which only have the Embedding transformer and thus cannot incorporate the *cis*-regulatory information between the core promoter and pCREs. The resulting performances were compared with the intact Chromoformer model. (c) Comparison of the performances for a subset of genes without or with known chromatin interactions. ROC-AUC scores of Embedding transformer-only Chromoformer and intact Chromoformer were computed only for a subset of genes that does not have known *cis*-regulatory interactions (Upper), and genes with at least one known *cis*-regulatory interactions. *p < 0.05, **p < 0.01 from two-sided paired t-tests.

We next examined whether the incorporation of distal *cis*-regulatory interactions into modeling truly contributed to the performance improvement. To this end, we eliminated Pairwise Interaction and Regulation transformers from the Chromoformer model to see how well it performs when trained without using distal *cis*-regulatory interactions. We retrained the ablated Chromoformer models and compared their performances to the intact Chromoformer model in terms of ROC-AUC (Figure 3b). This revealed that the Chromoformer in its intact form showed significantly high performances in most of the cell types analyzed (10 out of 11), implying that the inclusion of distal pCREs and their interactions into deep modeling helped learning epigenetic factors that govern the expression of genes. To further support that this improvement is specifically due to the modeling of *cis*-regulatory interactions, but not due to the mere complexity of the model, we investigated the modeling performance separately for the genes whose promoters do not interact with any pCREs. Since no biologically meaningful information would be transferred to the core promoter embeddings of those genes through the Pairwise Interaction and Regulation transformers, we can discern whether the dominant factor that contributed to the performance increase was the information transfer between genomic regions or the increased model complexity itself. As expected, we observed that Chromoformer showed no significant improvements for the majority of cell types (Figure 3c). Besides, the performance increase was still significant for the genes having at least one interaction with pCREs (Figure 3c). Together, it indicates that the contribution of *cis*-regulatory modeling is far greater than that of increased modeling capacity arising from deeper layers.

### Chromoformer learns to attend to the distant transcriptional elongation signals at gene bodies

Given the outstanding performance of the Chromoformer model compared to the other deep learning models, as well as the success of the wide-range modeling of core promoter region, we then asked if there are any unique patterns of HMs captured by Embedding transformers. As the self-attention layers inside the embedding transformers were designed to comprehend the dependencies of HMs at core promoters, we postulated that any such dependencies that the Chromoformer model is aware of can be revealed through the self-attention map it yields. Therefore, the attention weights produced by the Embedding transformer during the prediction were visualized to analyze the internal behavior of the model. Fig. 4a shows an example snapshot of attention weights during the prediction of the expression of anti-silencing function 1A (*ASF1A*) histone chaperone in the H1 cells (E003). Interestingly, we observed that a majority of attention heads were prominently giving strong attention to 4-6kbp downstream the TSS than any other specific positions (Figure 4a and b). This was rather unexpected since most of the regulatory information delivered by HMs are known to be deposited near the TSS, where the binding of transcription factors and transcriptional initiation mostly take place. In line with this notion, average signals of HMs displayed their characteristic patterns mostly at the TSS (Figure 4c). Specifically, methylations at H3K4 and histone acetylations were associated with higher levels of gene expressions, whereas high levels of H3K9me3 and H3K27me3 modifications were associated with low expression of genes.

**Figure 4.**
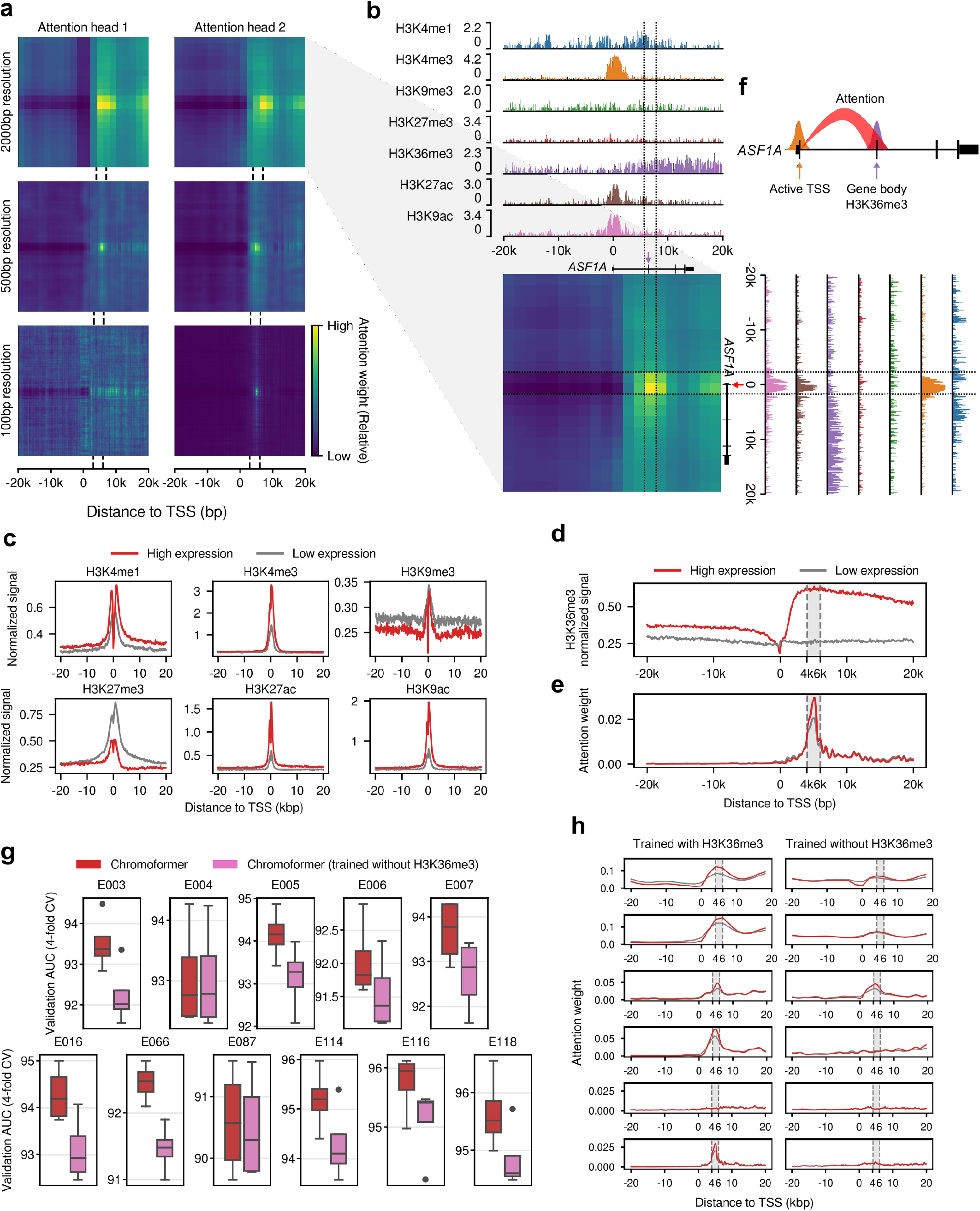
Analysis of self-attention weights learned by the Embedding transformer. (a) Representative self-attention weight matrices for the prediction of the expression of ASF1A in H1 cell. Each heatmap shows the attention weight for each pair of genomic bins. The dotted lines indicate the 4-6kbp region downstream TSS. (b) Detailed description of the attention weights learned by the attention head 2 for 2kbp-resolution Embedding transformer. Genome tracks representing the normalized signals of the seven HMs are aligned with the attention weight matrix. The dotted lines demarcate the regions 4-6kbp downstream the TSS. The red arrow indicates the TSS, and the purple arrow indicates the exon located within the region 4-6kbp downstream the TSS. (c) Average signals of the HMs other than H3K36me3. Signals were separately averaged according to their expression labels (High/Low expression). (d) Average H3K36me3 signal. (e) Average attention weights of the second attention head for 100bp resolution bins. The grey shade denotes the 4-6kbp region downstream the TSS. (f) Schematic diagram illustrating the behavior of the Embedding transformer. (g) H3K36me3 feature ablation experiment results. (h) Attention weights for TSS bin.

In striking contrast, the average H3K36me3 signal of the two classes of genes (i.e., high/low expression) did not show any notable difference at the TSS, but the difference was maximized at the 4-6kbp region downstream the TSS (Figure 4d). H3K36me3 is established by addition of methyl groups at H3K36 by SETD2 histone methyltransferase, and SETD2 is known to be recruited to the C-terminal domains of RNA polymerases in concert with the transcriptional elongation^22^. Thus, H3K36me3 is widely appreciated as a transcriptional elongation-associated HM that predominantly marks the bodies of actively-transcribed genes. The average attention weights imposed by the TSS-containing bin reached its maximum exactly at 4-6kbp region downstream the TSS (Figure 4e), suggesting that the model was well optimized to focus on the most discriminative genomic region in terms of H3K36me3 signals. Intriguingly, the extent of the attention given at the 4-6kbp region downstream TSS was far greater for genes with high expression than low expression (Figure 4e). In other words, the Embedding transformer was trained to adaptively control the amount of attention given on the HMs at the gene body, based on the histone context at the direct vicinity of the TSS, as illustrated in Figure 4f. One explanation for this behavior of the Embedding transformer is that the model seeks for the complementary evidence that reinforces its confidence for the initial guess on the gene expression, which is based on the HM states near the TSS. In this sense, H3K36me3 is a well-suited candidate for such a role since its discriminative power resides in where the other HMs do not show large variabilities, hence being left as the only clear signal in those regions. The importance of H3K36me3 at downstream gene bodies was further supported by the feature ablation experiment. When the H3K36me3 signals were excluded from model training and only the other six HMs were used as input features, we observed a significant decrease in performance (Figure 4g). Moreover, we discovered that the Embedding transformer predominantly lost the specific attention to the 3-6kbp downstream region (Figure 4h).

It seems that the importance of the feature representing transcriptional elongation was particularly important in this problem setting because the model was trained to predict the steady-state levels of mRNA measured by RNA-seq. Steady-state levels of mRNA are determined not only by the transcriptional initiation, but also by various factors including the rate of transcriptional elongation and mRNA stability. According to a study comparing different measurement technologies used for gene expression prediction tasks^23^, predicting the expression levels measured by cap analysis of gene expression (CAGE) was shown to be easier than predicting RNA-seq-based expression levels. This study also showed that H3K36me3 was predictive for RNA-seq-based expression levels, while core promoter HMs including H3K4me3 were more useful for CAGE measurements. They imply that the hidden factors including the efficacy of transcriptional elongation reside in RNA-seq measurements and they cannot be readily accounted for only with core promoter features. Thus, we speculate that the superior performance of Chromoformer model may arise from the ability to model the rate of transcriptional elongation, which leaves its trace as H3K36me3 in the gene body. These results, based on the great interpretability of the Embedding transformer, collectively suggest that the Embedding transformer learned the distant correlation between histone codes dictating active transcription near TSS and high levels of H3K36me3 representing transcriptional elongation at gene body, especially at the gene bodies 4-6kbp downstream the TSS. Moreover, this in part explains why the performance of Chromoformer showed consistent increase along the increase of window size, as it could collect additional evidence for gene expression also from the gene body.

### Chromoformer learns the additive transcriptional activation at transcription factories and switch-like suppression at PRC2-bound silencing hubs

We then examined the effect of modeling *cis*-regulation by distal pCREs in detail. As it has been already shown that the inclusion of Pairwise Interaction and Regulation transformers results in better overall performance, we sought for a more detailed interpretation for the gene-level effect of regulatory interaction modeling. To this end, we devised a gene-level measure to quantify the predicted impact of the *cis*-regulation based on the latent representation learned by Chromoformer. Specifically, we measured the Euclidean distance between the interaction-aware regulatory embedding produced by Pairwise Interaction and Regulation transformers, and the original core promoter embedding resulting from Embedding transformers (Figure 5a). For convenience, we termed this quantity as ‘predicted *cis*-regulatory impact (PCRI)’.

**Figure 5.**
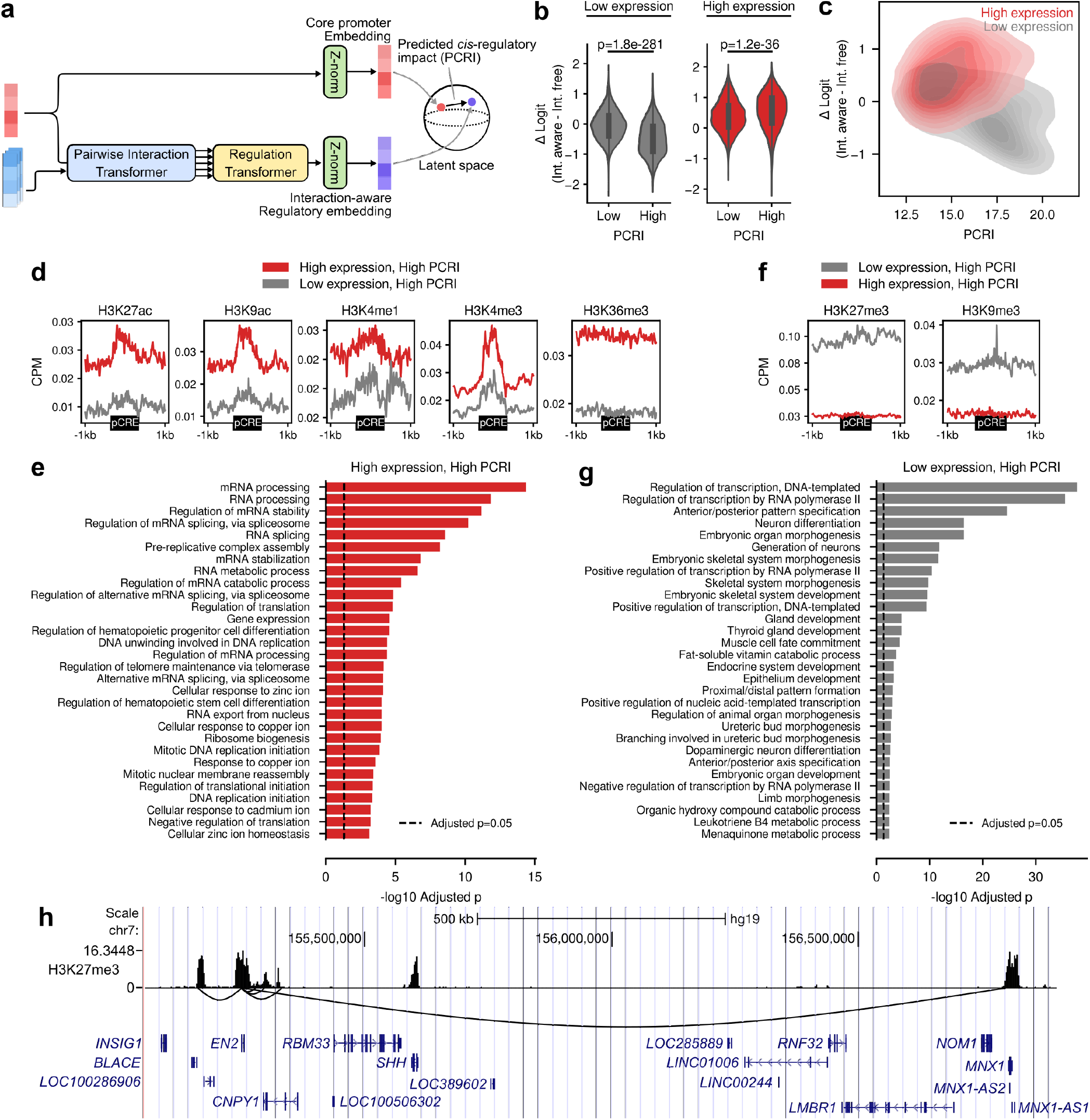
Analysis of predicted *cis*-regulatory impact (PCRI). (a) Schematic diagram illustrating the computation of the PCRI. (b, c) Relationship between PCRI and the difference of predicted logit (ΔLogit) between the interaction-aware and interaction-free Chromoformer. Shown are results for H1 cells (Reference epigenome ID E003). (d) Average HM signals near the pCREs interacting with top 1,000 genes with the highest PCRI among highly expressed genes. (e) Functional enrichment analysis of the top 1,000 genes with the highest PCRI among highly expressed genes. (f) Average HM signals near the pCREs interacting with top 1,000 genes with the highest PCRI among lowly expressed genes. (g) Functional enrichment analysis of the top 1,000 genes with the highest PCRI among lowly expressed genes. (h) Representative genomic region showing suppressive *cis*-regulatory interactions for *EN2*. Black curved lines below the H3K27me3 signal track shows the 3D chromatin interactions centered at the core promoter of *EN2*. NCBI RefSeq gene annotations are shown at the bottom.

To assure that PCRI truly reflects the dynamics of latent vector representations as well as the effect of regulatory interactions, we first asked how it eventually affected the prediction outcome probabilities. The difference in predicted logit (ΔLogit) between the interaction-aware Chromoformer and interaction-free Chromoformer was measured for each gene to determine the amount of perturbation on the prediction probability. As a result, ΔLogits for the two groups of genes with above and below half PCRI were significantly different within each of the low expression and high expression groups of genes in H1 cells (E003) (Figure 5b). Similarly, PCRI and ΔLogit for the highly expressed genes were positively correlated (Spearman’s r=0.15, p=6.04 × 10^−46^) and PCRI and ΔLogit for the lowly expressed genes were negatively correlated (Spearman’s r=-0.42, p<10^−308^) (Figure 5c). In short, it can be summarized that high PCRI for highly-expressed genes (labeled as 1) made those genes predicted with higher probability of gene expression (ΔLogit > 0), and high PCRI for lowly-expressed genes resulted in lower prediction probability (ΔLogit < 0). These results show that the regulatory impact of the pCREs on the core promoter was correctly captured by the Pairwise Interaction and Regulation transformer in terms of PCRI, hence it positively contributed to the prediction of gene expression states. They also suggest that those modules had learned to distinguish at least two types of *cis*-regulations, namely activation and suppression, based on the histone codes presented at pCREs.

To get deeper insights into how the Chromoformer model could accurately discern pCREs associated with the activation or suppression of gene expressions, we analyzed the characteristics of pCREs assigned for genes at the highest extreme in terms of PCRI, i.e., genes predicted to have greatest impact by distal *cis*-regulation. We collected 250 highly-expressed genes with the highest PCRI values from each fold of the 4-fold CV and examined the average signals of HMs near the pCREs associated with those 1,000 genes. As a result, pCREs for highly-expressed genes with high PCRI on average showed increased levels of HMs associated with transcriptional activation compared to those for lowly-expressed genes (Figure 5d). In particular, HMs representing enhancers (H3K27ac, H3K9ac, and H3K4me1), active promoters (H3K4me3) and active gene bodies (H3K36me3) were enriched for those pCREs. This broad enrichment of genomic elements associated with the greatest transcriptional activation implies that Chromoformer learned the existence of ‘transcription factories’, on which the active genes and enhancers are gathered together for efficient transcription^24^. Based on this observation, we sought for additional biological evidence by examining whether those genes clustered at putative transcription factories show enrichment for particular biological functions. Interestingly, they were highly enriched for housekeeping activities, including mRNA splicing, DNA replication, ribosome biogenesis and DNA damage response (Figure 5e). We also observed the enrichment of cell type-specific functions such as telomere maintenance in stem cells (E003 and E016), cell morphogenesis and extracellular structure organization in mesenchymal stem cells (E006) or iron homeostasis in liver cells (E066) or hepatocellular carcinoma (E118) (Supplementary Figure 3a). Taken together, it can be speculated that Chromoformer reflected the tendency of cells to ensure the robust expression of essential genes for its function and survival through sequestering them within transcriptionally active subcompartments harboring multiple enhancers^25^. On the other hand, pCREs for lowly-expressed genes with high PCRI on average showed increased levels of repressive marks such as H3K27me3 and H3K9me3 (Figure 5f), implying that Chromoformer also detected the transfer of the suppressive regulatory information from the pCREs to the core promoter. We conjectured that those pCREs represent transcriptional silencers, since previous studies have shown the potential functionality of distal H3K27me3-rich regions in transcriptional repression^26^. The top 1,000 genes which are predicted to have strong suppressive *cis*-regulation showed extreme enrichment towards developmental functions (Figure 5g). One representative example of the suppressive *cis*-regulatory interactions are shown in (Figure 5h) for Engrailed Homeobox 2 (*EN2*). As expected, many of the pCREs showed high H3K27me3 signals. Notably, one of the pCREs was located 1.5Mbp away from *EN2*, and the pCRE spanned the core promoter of Motor Neuron And Pancreas Homeobox 1 (*MNX1*), which is another homeobox transcription factor associated with development. The functional similarity between the two extremely distant, but interacting genes implies the existence of ‘silencing hubs’, where the developmental genes and silencers are sequestered together through 3D chromatin folding.

To assess the quantitative characteristics of the two regulatory hubs, we analyzed the association between the levels of PCRIs and the number of pCREs associated with transcriptional activation or suppression. First, to define transcriptionally active regions across the genome, we utilized the chromHMM-based chromatin state annotations available on the Roadmap Epigenomics Project. Among the 18 states, genomic intervals representing active TSS, active gene bodies or enhancers were considered as active regions, and we counted for each gene the number of pCREs overlapping with the identified active regions. We found that the number of the transcriptionally active pCREs and PCRI showed a moderate but significant positive correlation (Spearman’s r=0.13, p=5.2 × 10^−34^) (Figure 6a). As the expression levels of genes also increased in accordance with the increasing number of transcriptionally active pCREs (Figure 6b), it implies that Chromoformer learned the additive dynamics of pCREs in transcription factories for gene expression activation.

**Figure 6.**
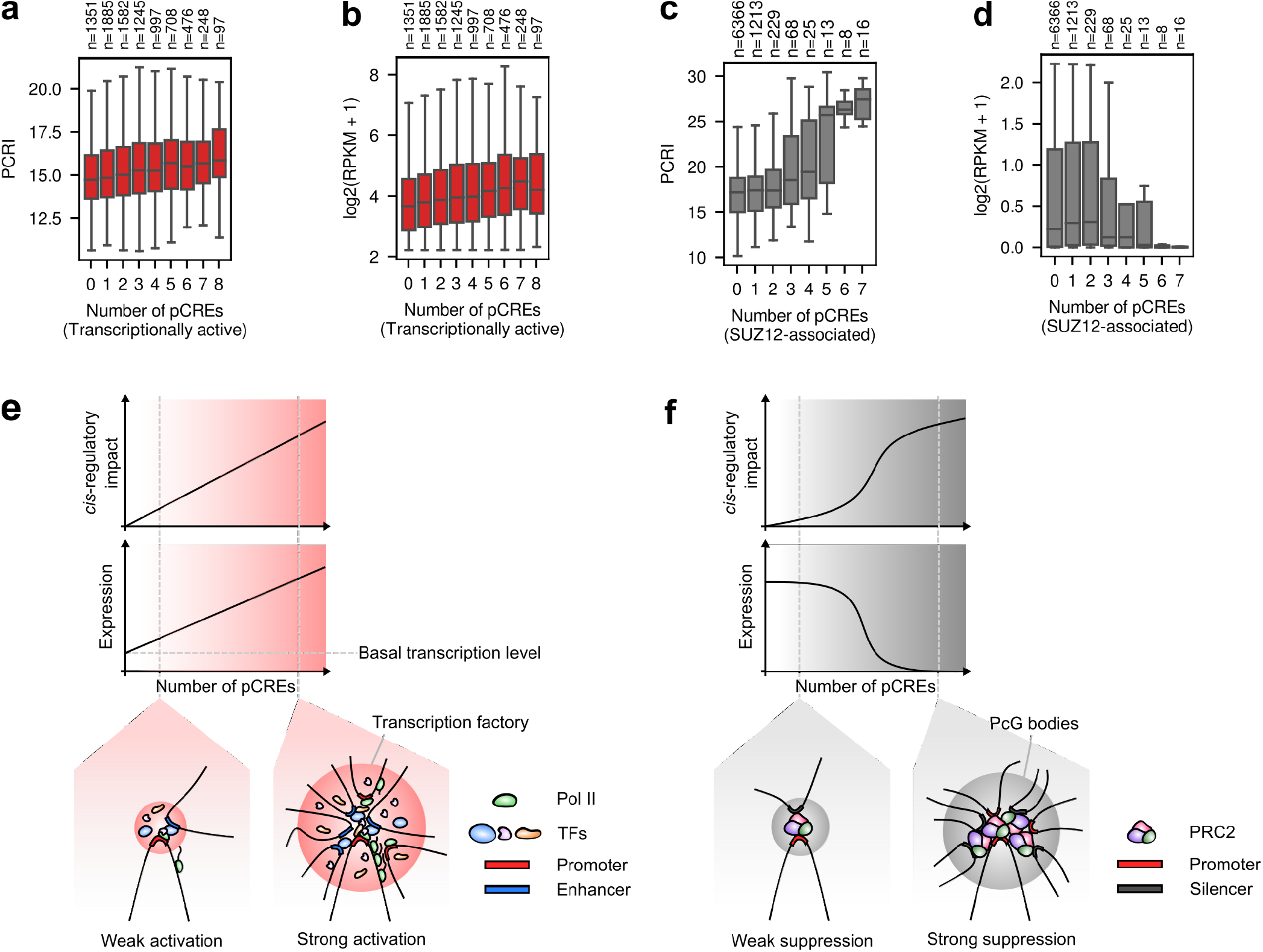
Characteristics of *cis*-regulation learned by Chromoformer. (a) Association between the number of transcriptionally active pCREs and PCRI. (b) Association between the number of transcriptionally active pCREs and gene expression level. (c) Association between the number of pCREs harboring SUZ12 binding sites and PCRI. (d) Association between the number of pCREs harboring SUZ12 binding sites and gene expression level. Illustrations for the proposed hypothetical models for regulatory dynamics of (e) transcription factories and (f) silencing hubs are shown. Pol II, RNA polymerase II; PcG bodies, Polycomb group bodies; PRC2, Polycomb repressive complex 2.

For silencing hubs, we considered the pCREs associated with Polycomb group (PcG) proteins as functional silencers. Beside its enzymatic role as a histone methyltransferase targeting H3K27, it has been recently demonstrated that Polycomb repressive complex 2 (PRC2) functions as a mediator of the repressive *cis*-regulatory interaction between silencers and developmental promoters, promoting the formation of PcG bodies^27^. To determine the number of PRC2-bound silencers, we utilized the ChIP-seq peaks for the two of the three core subunits of PRC2, namely SUZ12 and EZH2, from ENCODE. As a result, we found the logistic increase of PCRI upon the increasing number of pCREs overlapping with the SUZ12 binding site (Figure 6c), which indicates that the switch-like transcriptional suppression by PRC2-mediated silencing hub was learned by Chromoformer. Similar results could be reproduced for EZH2 binding (Supplementary Figure 3b and c). Notably, this trend entirely disappeared when tested for the number of non-specific pCREs regardless of the association with PRC2 (Supplementary Figure 3d), which is suggestive of the highly specific nature of PcG-mediated silencing.

To date, the collective effect of distal *cis*-regulatory elements on gene expression remains incompletely understood, but nevertheless, the pioneering works exploiting modern technologies such as STARR-seq^28^ or CRISPRi-FlowFISH^29^ certainly provide us with deep insights about their dynamics. Intriguingly, the observations drawn from the interpretation of trained Chromoformer models, which are optimized to capture the quantitative characteristics of *cis*-regulation, highly agree with the latest viewpoints from such studies. Our observations on the additive transcriptional activation by active pCREs recapitulates the results of a previous study on the quantitative characterization of enhancer activity in *Drosophila*. The underlying mechanism for this additivity has been explained by either of the ‘interaction hub’ or ‘promoter competition’ model^30^. The former assumes a multi-way interactions between a promoter and several enhancers with independent contributions, while the latter posits the one-to-one promoter-enhancer interactions and demonstrates that the probability of the contact between a promoter and any enhancer increases as the number of candidate enhancers increases. On the contrary, the quantitative nature of transcriptional silencing by PcG bodies with regard to the number of PcG-bound silencers is yet to be fully characterized. Our interpretation of Chromoformer leads to the hypothesis that there exists certain threshold of the local concentration of silencers for PcG bodies to fully exert their suppressive function (Figure 6c). It may be due to synergy with other repressive epigenetic factors including the DNA methylation induced by the HMs newly added by those PcGs and other chromatin remodeling factors. In any case, experimental validation of this hypothesis and further characterization of the biological factors that determine the tipping point of the PcG-mediated gene silencing will highly improve our understanding of precise regulation of gene expression. Altogether, these results demonstrate the utility of Chromoformer and, by extension, deep learning models in the derivation of the new quantitative hypothesis in the field of computational biology that would ultimately facilitate experimental validations and thus new scientific discoveries.

## Discussion

In the present study, we proposed a novel transformer-based deep learning architecture named Chromoformer to model the quantitative role of the histone codes in the regulation of gene expression. Chromoformer greatly improved the performance of gene expression prediction by modeling the three-level hierarchy of *cis*-regulation involving core promoters and pCREs. By the analyses of self-attention weights and latent embedding dynamics and several ablation studies, we also provided in-depth biological interpretations regarding the behavior of the Chromoformer model. Thanks to the power of transformers for comprehending distant dependencies in a sequence, Chromoformer could successfully learn to focus on the specific region inside gene bodies where the HMs associated with gene expression were the most distinctive between highly expressed and lowly expressed genes. Interestingly, the amount of attention paid to the gene body was dependent on the epigenetic context of the TSS, implying that the Chromoformer model captured the distant dependencies of the HMs placed at TSS and gene body. On the other hand, by using transformers to model pairwise relationships within an unordered set of features, Chromoformer could learn how the information mediated by histone code is propagated from pCREs to core promoters through 3D chromatin folding to regulate gene expression. Analysis of the latent representations of histone codes learned by the model highlighted that the expression of housekeeping and cell-type specific genes were reinforced by the interaction with enhancers, whereas the expression of developmental genes were mainly repressed by the interaction with PRC2-bound silencers.

We explicitly used a pre-compiled knowledge of 3D chromatin interactions to guide Chromoformer learning. Those experimentally measured interaction frequencies were used to prioritize the pCREs that will participate in the model training by being explicitly injected into the self-attention score matrices. However, it also seems possible to infer the interaction frequencies between pCREs and the core promoters from genomic sequence information alone. This is because the specificity of *cis*-regulatory interactions are largely governed by the recognition of DNA sequence motifs by DNA-binding proteins including transcription factors or CCCTC-binding factors (CTCFs), which function as insulators that compartmentalize the 3D genome conformations. Therefore, those binding motifs embedded in the genome may serve as hidden vocabularies that allow the inference of the desired chromatin conformations solely based on the DNA sequence. Results from the recent model named Enformer strongly supports that such *de novo* prioritization of pCREs are more effective when wider sequence information is used^31^, thereby suggesting the exciting possibilities for the fully data-driven modeling of gene expression regulation through the integration of genomic and epigenomic features using the transformer architecture. We leave this transformer-based multi-omics integration as a further work.

The attention learned by the Embedding transformer that jumps from an active TSS to the gene body suggests that the HMs placed at gene bodies are indeed useful, if not the most critical, information when predicting the steady-state gene expression levels. From this result, we can consider the possibility that using the entire landscape of histone codes distributed throughout a single gene may further improve the predictive accuracy for steady-state mRNA levels. Furthermore, as H3K36me3 is far more enriched at exons than introns, utilizing the full-length gene annotation will be another effective guidance for model training. As gene lengths and exon-intron distributions show great variability, we need some clever representation of such biological prior knowledge. Again, the transformer architecture would be one of the most powerful choices because one can flexibly apply masks to deal with variable-length inputs and also can extend positional encoding to form composite encoding that simultaneously harbors information for both genomic positions and annotations for gene structures.

In summary, Chromoformer is another exemplary application emphasizing the great potential of transformer architecture in modeling of biological sequences. This study also underscores the importance of developing specialized deep learning architectures effectively embedded with biological prior knowledge, not only for the improvement of the performance in predictive tasks, but also quantitatively characterizing the complex relationships between biological entities.

## Methods

### Promoter-centered 3D chromatin interactions

Experimentally validated core promoter-pCRE interaction pair information is required to train a Chromoformer model. In this study, we used publicly available data deposited in the 3DIV database for promoter-centered long-range chromatin interactions compiled by pcHi-C experiments comprehensively conducted for various tissue types. Interactions were characterized at the resolution of HindIII restriction fragments with median and average length of 4,797bp and 5,640bp, respectively. We could also obtain normalized interaction frequencies between the DNA fragments. These frequencies were normalized by a two-step normalization that accounts for the capturability and distance-dependent background signals. Although the significant interactions could be selected based on the estimated FDR values provided along with each interaction, we considered an interaction as significant if the normalized frequency was greater than 1.5 to increase the sensitivity of chromatin interactions during the Chromoformer training. Note that a normalized frequency of 1.5 denotes that the ratio between the interaction and background signal is 1.5.

### Training data preparation

Consolidated ChIP-seq read alignments for seven major HMs (H3K4me1, H3K4me3, H3K9me3, H3K27me3, H3K36me3, H3K27ac and H3K9ac) were obtained from the Roadmap Epigenomics Project^19^. This alignment data could make a highly homogeneous training dataset, since the reads were truncated to 36bp to reduce read length bias originating from the difference in sequencing experiments, and also they were subsampled to a maximum of 30 million reads to homogenize for read depths. Read depths along the hg19 reference genome were computed for each base position using ‘genomecov’ command of Bedtools^32^. For each given core promoter or pCREs, we computed log2-transformed binned average signals of seven HMs along non-overlapping genomic windows of size 100bp, 500bp and 2000bp that fully covers the region and used those values as input features for our model. Since pCREs have different sizes in our setting, we zero-padded pCRE feature matrices to make their size agree with that of the core promoter feature matrices. Specifically, we center-aligned the matrix and appended zero-matrices of appropriate size to the left and right side of the input matrix. To determine prediction target labels, normalized gene expression levels (in RPKM) were also downloaded from the Roadmap Epigenomics Project. RefSeq annotation was used to determine the TSS for each gene. Total 18,955 genes that were appropriately annotated and had expression measurements were selected for model training and evaluation. For each cell type, the median expression values across all genes were used as threshold values to assign genes with one of the two labels: highly (1) or lowly expressed (0). This formulation of binary classification of gene expression has been widely adopted for various machine learning approaches for gene expression modeling.

### Selection of cell types for model training

For model training, we only chose a subset of cell types analyzed in the Roadmap Epigenomics Project for which the whole profiles of gene expression, HMs and 3D chromatin interactions were available. Since 3D chromatin interaction profiles are not the official results of Roadmap Epigenomics but are obtained from an independent source, we manually matched the EIDs and cell type mnemonics from 3DIV database^17^. As a result, the following 11 cell types were used for Chromoformer training: H1 cells (E003, H1), H1 BMP4 derived mesendoderm (E004, ME), H1 BMP4 derived trophoblast (E005, TB), H1 derived mesenchymal stem cells (E006, MSC), H1 derived neuronal progenitor cultured cells (E007, NPC), HUES64 cells (E016, H1), Liver (E066, LI11), Pancreatic islets (E087, PA), A549 EtOH 0.02pct lung carcinoma (E114, LG), GM12878 lymphoblastoid (E116, GM) and HepG2 hepatocellular carcinoma (E118, LI11).

### Chromoformer model architecture

Chromoformer consists of three transformer-based modules, Embedding, Pairwise Interaction and Regulation transformer.

#### Embedding transformer

Embedding transformer has a single encoder layer, which takes a binned average signal matrix *X*_input_ of seven HMs at a core promoter and summarizes it into a core promoter embedding matrix *X*_emb_ that consists of fixed-sized latent embedding vectors. Before *X*_input_ is fed into the module, seven-dimensional input features for each of the *n* bins are first linearly projected into the dimension of *d*_emb_ (= 128), then a positional encoding matrix *P* of dimension *n× d*_emb_ is added to the input feature matrix of the same dimension in an element-wise manner. *P*_*ij*_ is defined as below.

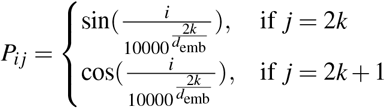

It is worth noting that the inner product between any two row vectors in the positional encoding matrix *P*, namely *P*_*a*_ and *P*_*b*_, only depends on the positional distance |*a− b*| between the two vectors. Therefore, by adding a positional encoding matrix to the input feature matrix, the relative distance between any two features can be recognized in the following multi-head attention layers. A multi-head attention layer in Embedding transformer utilizes a self-attention mechanism to capture inter-dependencies between HMs that contribute to the regulation of gene expression. Importantly, those operations are done separately for multiple heads so that the model can capture different aspects of inter-dependencies between input features. Self-attention operation in the transformer architecture is a special case of scaled dot-product attention where the query, key and value matrices originate from the same sequence of features. Specifically, position-encoded input feature matrix of dimension *n × d*_emb_ is linearly projected to produce three matrix *Q*_emb_, *K*_emb_ and *V*_emb_ of dimension 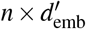, which semantically represents a query, key and value matrix, respectively. 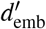 is set to 64. The *n × n* matrix produced by a multiplication of *Q*_emb_ and 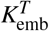 is called a pairwise affinity matrix since each element of the matrix is equivalent to a dot product between the corresponding pair of vectors from *Q*_emb_ and 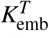 and denotes an amount of affinity between the two positions in the input sequence. The pairwise affinity matrix is divided by 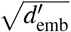 and softmax function is applied to convert self-attention affinities into a weight that sums to 1 for each row. The value matrix *V*_emb_ is multiplied with the resulting attention weight matrix to finally produce the output of self-attention operation. The whole process of scaled dot-product can be summarized as below:

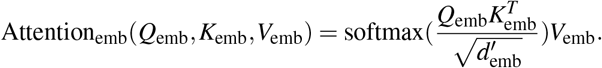

In the Embedding transformer, the self-attention is separately done by *m*_emb_ (= 2) heads and the resulting *m*_emb_ vectors of dimension 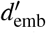 are concatenated to form a single *d*_emb_ (= 128) dimensional vector so that the dimension of input feature is preserved. Subsequently, the input sequence of features right before the self-attention is added via residual connection and then they are layer-normalized. The result is then subjected to the linear projection layer into the dimension of *δ*_emb_ (= 128), nonlinear activation by rectified linear unit (ReLU) and final linear projection into the dimension of *d*_emb_. This series of operations involving linear projection, nonlinear activation and another linear projection comprises a position-wise feedforward layer.

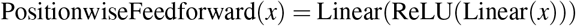

After another residual connection and layer normalization, the core promoter embedding matrix *X*_emb_ is finally produced.

#### Pairwise Interaction transformer

Pairwise Interaction transformer consists of two stacked layers to update a core promoter embedding based on its pairwise interaction with each pCRE and produce pairwise interaction embedding matrix *X*_pair_. The difference between encoder-decoder attention and self-attention operation in the Embedding transformer is that encoder-decoder attention separately builds query matrix and key-value matrices. Specifically, query matrix *Q*_pair_ is derived from from *X*_emb_ (or *X*_pair_ resulting from the first layer), while key and value matrices *K*_pair_ and *V*_pair_ are built from position-encoded pCRE HM features *X*_HM_. In short, the query, key and value matrices for the encoder-decoder attention can be summarized as below.

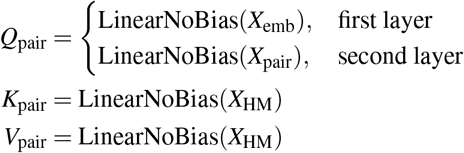

Then scaled dot-product attention is conducted between core promoter queries and pCRE key-values as below, where 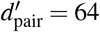 and *d*_pair_ = 128:

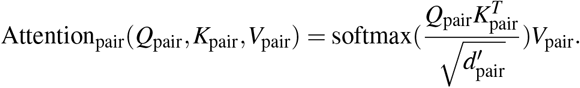

The rest of the operations including position-wise feedforward layer with *δ*_pair_(= 256), residual connections and layer normalizations are the same as the Embedding transformer, finally producing the pairwise interaction embedding *X*_pair_.

To avoid excessive computational load and to make the training batch fit in the memory of a single graphical processing unit (GPU) during training, we only considered at most *i*_max_ (= 8) pCREs for each core promoter. To determine the set of pCREs participating in the training, all the candidate pCREs were prioritized according to their normalized interaction frequencies with the core promoters since the pCRE that is interacting more frequently is likely to be more informative in predicting the expression of the corresponding gene.

#### Regulation transformer

Regulation transformer consists of six stacked layers with gated self-attention mechanism. The key function of the Regulation transformer is to update *X*_emb_ along with the whole set of *X*_pair_’s at the same time to finally produce the regulatory embedding *X*_reg_. To this end, individual embedding vectors that exactly represent the genomic bin where the relevant TSS is located are extracted from *X*_emb_ and *X*_pair_’s. Then, they are concatenated to form a composite input matrix *X*_comp_ of dimension (*i*_max_ + 1) *×d*_emb_ (Recall that *d*_emb_ = *d*_pair_ = 128). Specifically, those are the vectors at the midpoint of the embedding matrices. Note that for genes having less than *i*_max_ *cis*-regulatory interactions, the rest of *X*_comp_ was filled with dummy zero vectors.

The Regulation transformer does not need a positional encoding since it does not assume any predefined order among the embeddings. We only fix that the first row vector of the composite input matrix is the core promoter embedding. This unordered set of embeddings is fed to a gated self-attention mechanism to allow the model to decide how much it will actively utilize the transformed embedding carrying the interaction information. In addition to the query, key and value matrices, gated self-attention introduces a gate matrix *G*_reg_ that learns the amount of information transfer. The four 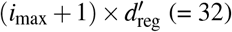 matrices used for gated self-attention operation are computed as below:

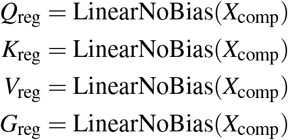

Moreover, we added a vector of normalized interaction frequencies *f* between the corresponding core promoter-pCRE pair as a bias term to the self-attention matrix to inform the model with the relative affinities of the pairwise interactions. An (*i*_max_ + 1) *×* (*i*_max_ + 1) bias matrix *B* is introduced, whose first row is filled with *f* and all the other values are zero. Taken together, the attention operation used in Regulation transformer can be written as below:

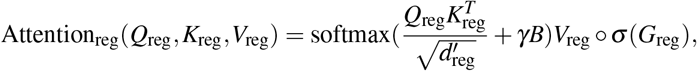

where *γ, σ* and ○ represent a learnable scalar coefficient, the sigmoid function and the Hadamard product, respectively.

We concatenated three *X*_reg_’s resulting from independent Chromoformer modules learning from 100bp, 500bp and 2000bp-resolution inputs, respectively. Only the first row of the concatenated matrix, which denotes the *cis*-regulation-aware embedding vector of the core promoter was extracted and fed into the fully-connected head. The fully-connected head has a single 128-dimension hidden layer and finally produces a two dimensional output representing the two prediction logits for each class.

### Model training and evaluation

Chromoformer was trained for 10 epochs with AdamW optimizer^33^ and the model resulting from the last epoch was chosen as the final model. The initial learning rate was chosen as 3 *×*10^−5^ and was decreased by 13% after each epoch so that it can approximately shrink to half of its value after each of the five epochs. Cross-entropy between the predicted probability and one-hot encoded binary gene expression label was used a loss function. Batch size was fixed to 64. All the implementations for benchmark deep learning models were obtained from the official code repositories provided by respective authors. To train the benchmark models, we applied the optimal hyperparameters that were previously identified for each benchmark model.

For GC-MERGE training, we needed to modify our input representations of HM signals and *cis*-regulatory interactions as per required. ChIP-seq read depths for each 10kbp bin throughout the genome were calculated using ‘multicov’ command of Bedtools, and the interaction frequencies between those 10kbp genomic bins were determined using the pcHi-C experiment results. Since GC-MERGE predictions are made for each of the 10kbp bins, but not for each gene, ambiguity arises when there are two or more genes in the same bin. The ambiguity is resolved by choosing a representative gene within each bin and assigning it with the most frequently occuring labels in that bin. This is done at the cost of reduced number of predictable genes. To perform as fair comparison as possible, we went through 4-fold CV for both GC-MERGE and Chromoformer using the same gene set whose expression is predictable by GC-MERGE, by retraining Chromoformer model.

### Analysis of Embedding transformer self-attention

Each Embedding transformer has two independent attention heads, so it produces two corresponding self-attention pairwise affinity matrices 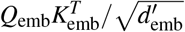 or each input. Since the full model consists of the three independent Chromoformers, we can extract six self-attention weight matrices in total. We visualized the softmax-normalized pairwise affinities, i.e., self-attention weights, for Figure 4. Note that all the self-attention weights were obtained at the time of inference for genes in validation set.

### Computation of the predicted *cis*-regulatory impact (PCRI)

To compute PCRI, we first defined a multi-resolution core promoter embedding as the concatenation of individual core promoter embeddings resulting from the three different input resolutions. Then, PCRI is defined as the Euclidean distance between the multi-resolution core promoter embedding and the multi-resolution regulatory embedding for each gene. Importantly, we standardized each embedding vector before calculating the Euclidean distance to correct for the global shift arising from the transformation itself. Similar to the self-attention analysis, the entire PCRI values discussed in the main text were calculated for genes in each validation set to ensure that the model did not explicitly learn the optimal transformation of latent embeddings reflecting *cis*-regulations for those genes.

### Functional enrichment analysis of genes with high PCRI

For each of the four validation folds, we identified top 250 genes with the highest PCRI values separately for each binary label. Functional enrichment analysis of the union of the four gene sets were done using Enrichr^34^. Gene ontology biological process terms with adjusted P-values < 0.05 were selected as significantly enriched terms.

## Author contributions statement

D.L. and J.Y. conceived the experiment(s), D.L. and J.Y. conducted the experiment(s), D.L., J.Y. and S.K. analysed the results. All authors reviewed the manuscript.

## Additional information

The source code for Chromoformer model are available at the following url: https://github.com/dohlee/chromoformer.

## Supplementary information

**Supplementary Figure 1.**
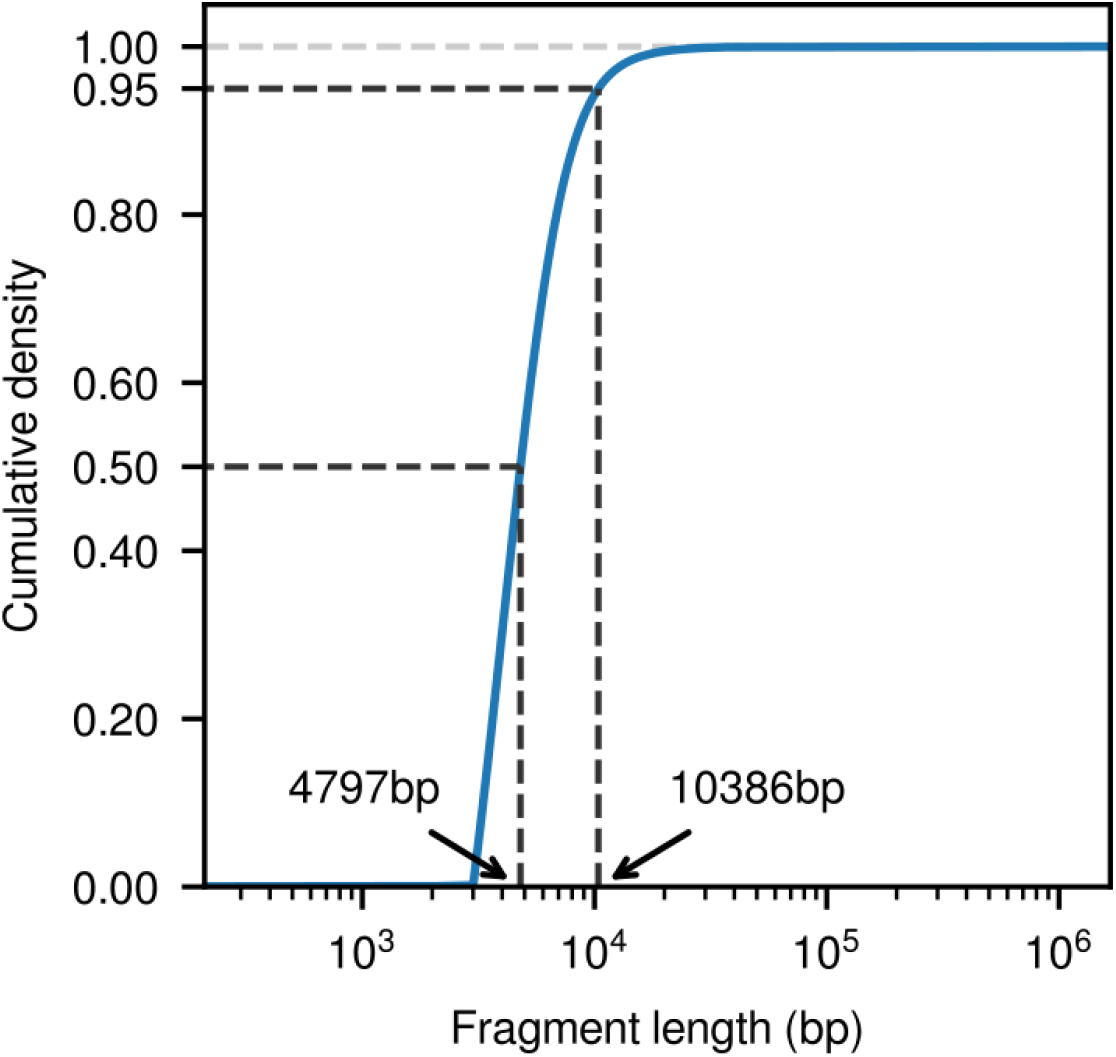
Distribution of HindIII fragment length from the pcHi-C dataset used in this study.

**Supplementary Figure 2.**
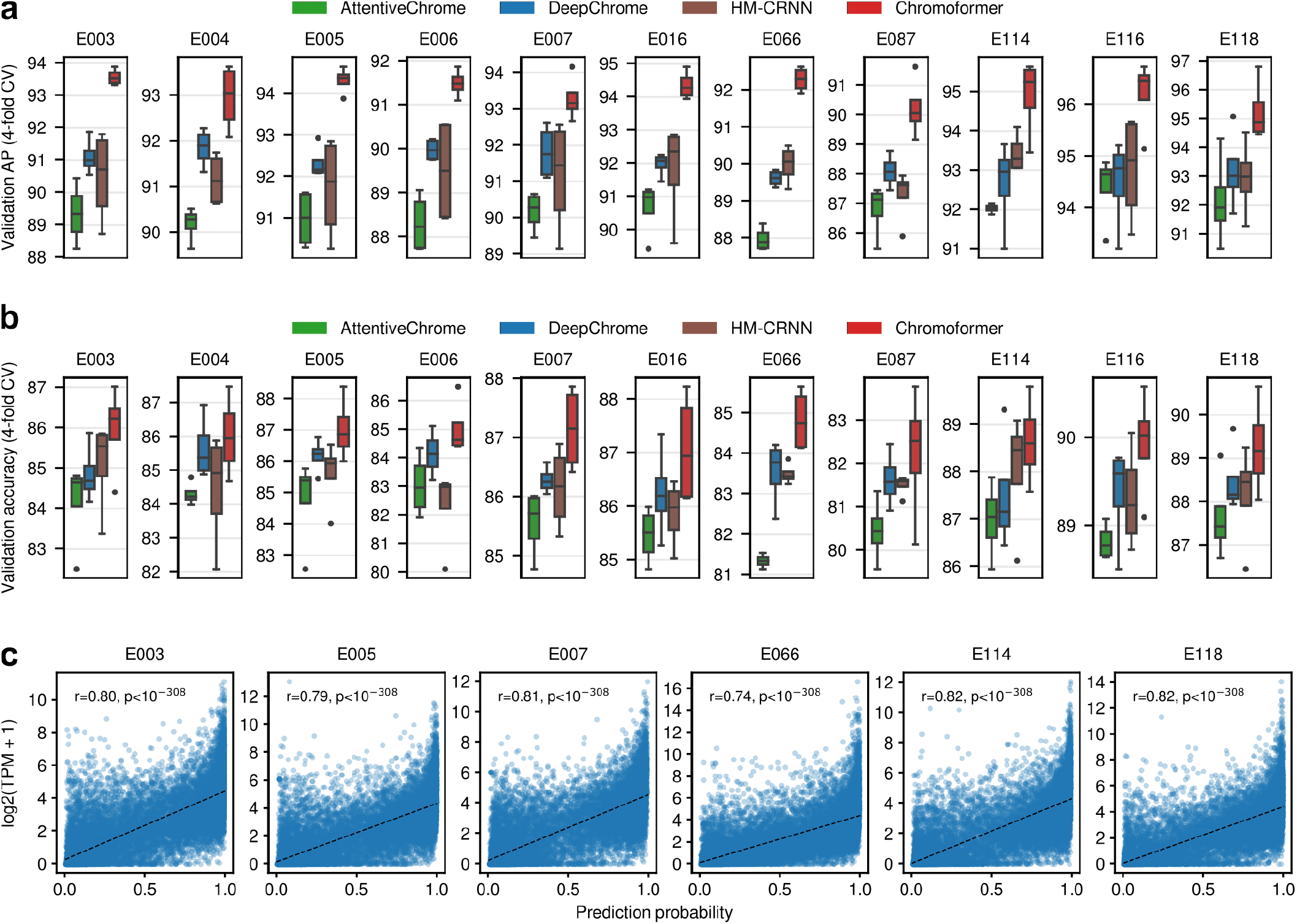
Comparison of (a) average precision and (b) accuracy between benchmark models and Chromoformer. (c) The prediction probability was highly correlated with the actual expression levels (Pearson’s r > 0.7, all p < 10^−308^). AP, average precision.

**Supplementary Figure 3.**
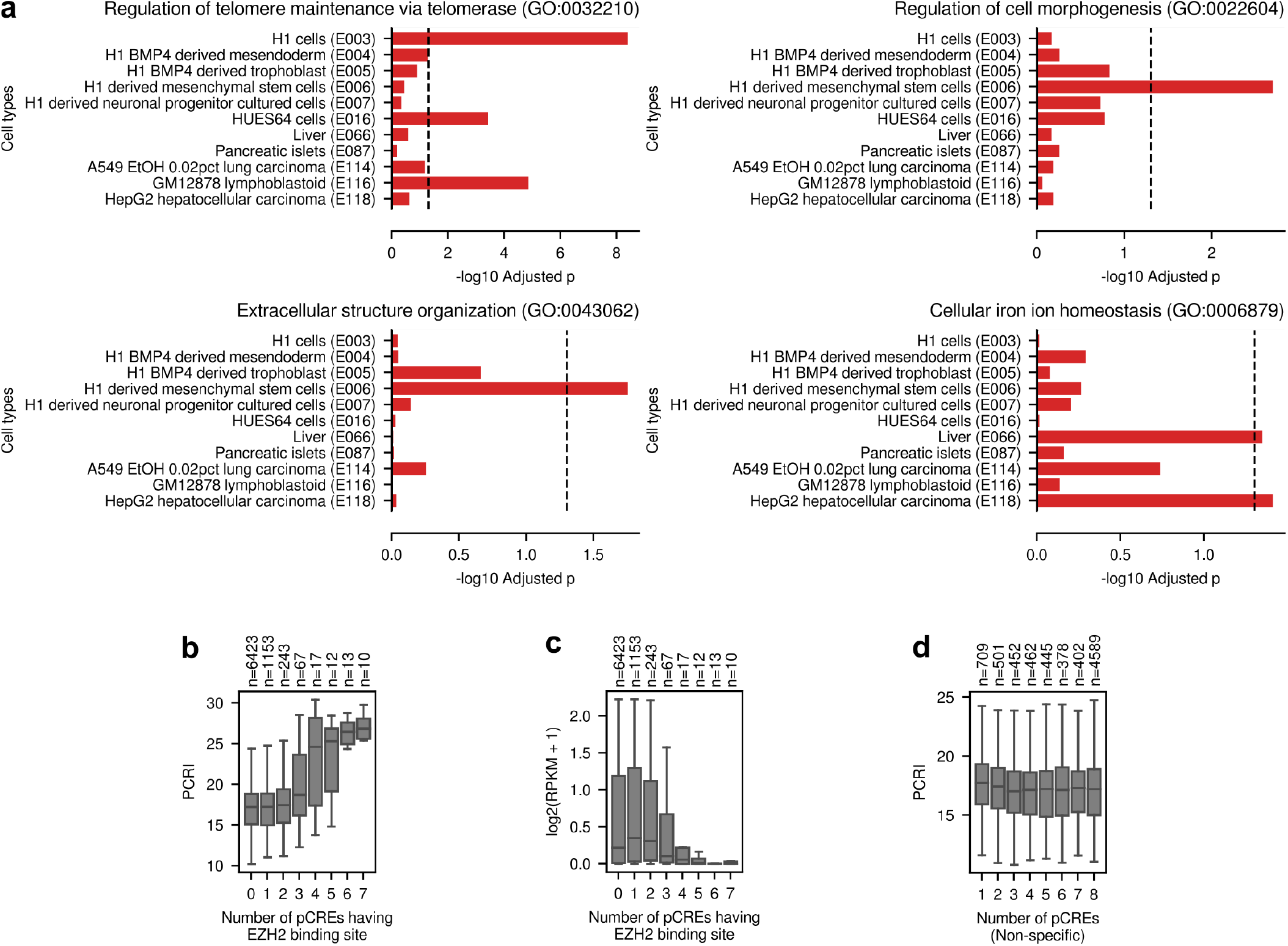
(a) Functional enrichment of highly expressed genes (i.e., labeled as 1) with high PCRI. For each cell type, top 250 genes with the highest PCRI values were selected for each of the four CV folds. Bars denote the adjusted p-values for the functional enrichment of the resulting 1,000 genes for each cell type. (b) The number of pCREs harboring EZH2 binding site versus PCRI. (c) The number of pCREs harboding EZH2 binding site versus the actual expression level of the corresponding gene. (d) The number of non-specific pCREs versus PCRI.

